# Resting-State Electroencephalography Alpha Dynamic Connectivity: Quantifying Brain Network State Evolution in Individuals with Psychosis

**DOI:** 10.1101/2024.06.04.597416

**Authors:** Romain Aubonnet, Mahmoud Hassan, Paolo Gargiulo, Stefano Seri, Giorgio Di Lorenzo

## Abstract

This study investigates brain dynamic connectivity patterns in psychosis and their relationship with psychopathological profile and cognitive functioning using a novel dynamic connectivity pipeline on resting-state EEG.

Data from seventy-eight individuals with first-episode psychosis (FEP) and sixty control subjects (CTR) were analyzed. Source estimation was performed using eLORETA, and connectivity matrices in the alpha band were computed with the weighted phase-lag index. A modified k-means algorithm was employed to cluster connectivity matrices into distinct brain network states (BNS), from which metrics were extracted.

The segmentation revealed five distinct BNSs. FEP exhibited significantly lower connectivity power in BNS 2 and 5 and a greater duration dispersion in BNS 1 than CTR. Negative correlations were identified between BNS metrics and negative symptoms in FEP. In CTR, correlations were found between BNS metrics and cognitive domains.

This analysis method highlights the variability of neural dynamics in psychosis and their relationship with negative symptoms.

## Introduction

Psychosis is a multifaceted mental condition characterized by a significant disruption in the perception and cognition of reality, often manifesting through delusions, hallucinations, disorganized thought and speech processes, and self-disturbances, as well as cognitive and negative symptoms, particularly in primary or idiopathic psychoses, such as schizophrenia ^1–3^. Functional neuroimaging research has consistently shown that psychotic disorders are associated with altered patterns of brain function, both at rest and during the execution of cognitive tasks (see ^4^ for a critical review).

Despite extensive research, inter-individual variability in patterns of brain function has limited the identification of a unifying model of neural mechanisms underlying psychosis ^5–8^, hindering the application of targeted diagnostic tools and early identification of the most effective treatment. This underscores the need for a new approach that can define patient-specific profiles.

Among the available functional neuroimaging techniques, electroencephalography (EEG) provides a unique opportunity to correlate recurring activity patterns with their underlying brain functional properties, contributing to describing what we now define as brain states ^9–11^. Its unsurpassed temporal resolution has proven to be a valuable tool in studying intermediate phenotypes in psychosis ^12–14^. Previous studies have reported abnormalities in brain electrical activity in psychotic disorders, such as decreased activity in the 8-12 Hz (alpha) frequency band ^15,16^. In addition to spectral EEG measures, EEG analysis techniques have increasingly focused on patterns of functional brain connectivity ^17^ and applied these methods to schizophrenia, identifying a disruption in the alpha-band connectivity ^18^. Magnetoencephalography (MEG) has confirmed network differences between psychosis and controls in the alpha band ^19,20^. Additionally, *Phalen et al.* ^19^ observed network features associated with psychopathological domains in psychosis.

Traditional static connectivity measures, often combined with graph theory metrics, may not adequately capture brain activity’s complex and rapidly changing nature ^21^. Measures of dynamic connectivity and new methods to model the transient states of brain networks have been developed to monitor and decipher these complex electrophysiological oscillations ^22,23^.

The current work introduces a novel methodology to analyze resting-state EEG (rsEEG) that integrates source localization techniques with dynamic functional connectivity and advanced clustering algorithms. This approach is inspired by the classical concept of EEG microstates, which have been used to capture the dynamic aspects of brain electrical activity ^24,25^. Defining the specific temporal properties of the EEG signal and connectivity patterns allows us to identify and characterize distinct brain network states (BNS). This method has recently been successfully applied to extract EEG dynamic connectivity patterns ^26,27^. The current study aimed to investigate source-space functional dynamic connectivity analysis in the rsEEG alpha band in individuals with first-episode psychosis (FEP) and healthy control subjects (CTR). Furthermore, we explored the potential relationships between these network (dynamic) states and measures of cognitive performance and psychopathological profiles. We hypothesize that BNS features may be related to cognition in both groups and, specifically, to negative symptoms in psychosis.

## Results

### 1. Clustering Results

After applying the modified k-means algorithm to the EEG connectivity matrices, we identified five distinct BNS. Notably, BNS 2, 4, and 5 were characterized by a greater strength of visual regions. Besides this visual strength, BNS 4 revealed a more substantial activity in ROIs within the left hemisphere. In contrast, BNS 5 showed more significant activity in the right hemisphere, particularly for DAN and SAN. BNS 2 shows a balanced and almost symmetric activity in both hemispheres, mostly DAN and SAN. A greater strength of the DMN nodes marked BNS 1 and 3. On top of its DMN prevalence, BNS1 has more present activity in the left hemisphere, especially in the SAN and MOT networks, whereas the DAN activity is symmetric. For BNS 3, we observe stronger activity in the “Other” RSNs in the right hemisphere but symmetric activity in the DAN and SAN. Figure 1 shows a representation of the BNS and their prevalent function.

**Figure 1.**
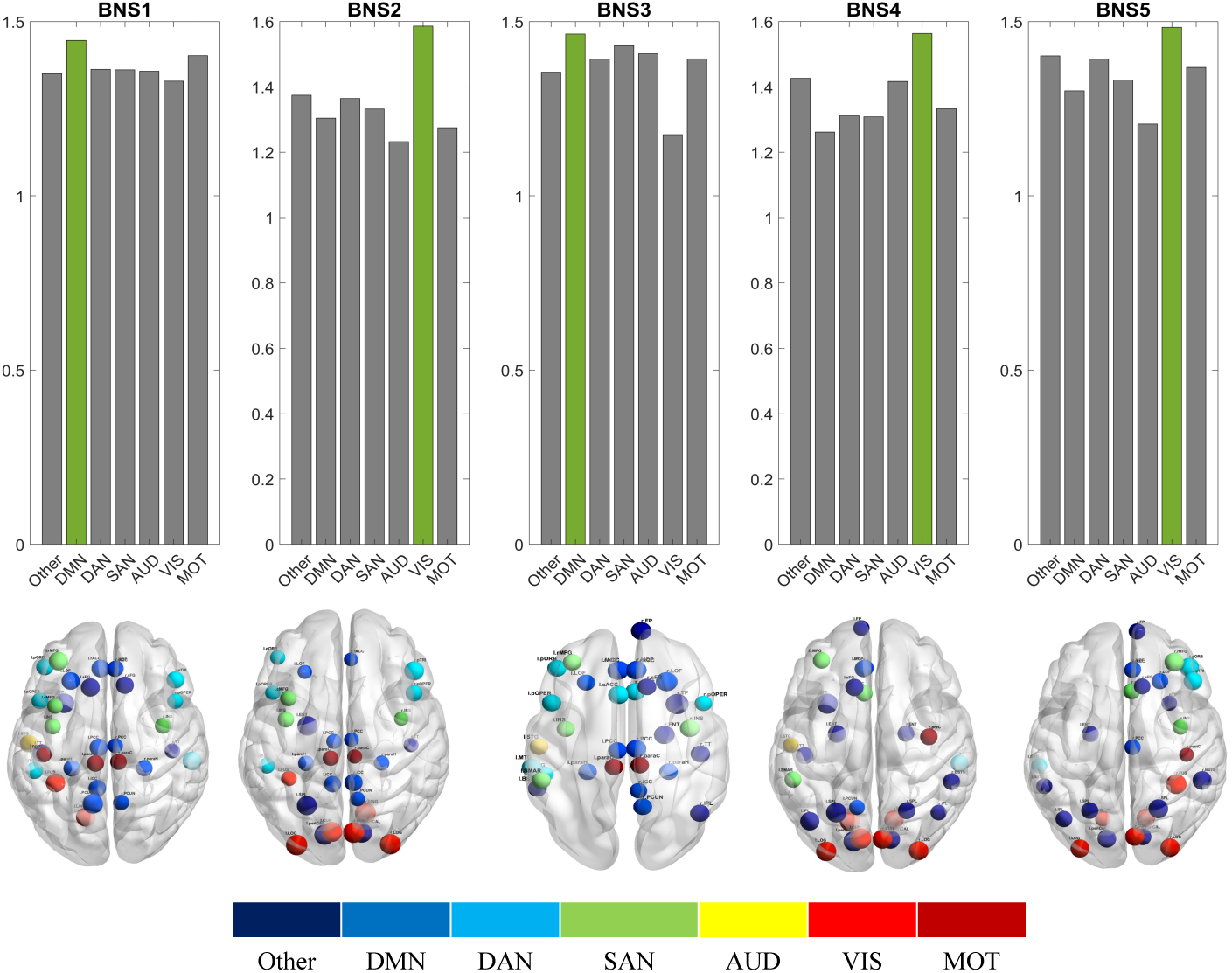
The resulting BNS with their prevalent function and node distribution based on strength.

### 2. Pairwise Comparisons

Upon comparing the BNS metrics between the FEP and CTR, significant differences emerged in the following metrics: GCP, standard deviation of duration, and transition probabilities, as highlighted in Figure 2. Specifically, BNS 2 and 5 exhibited a lower GCP in FEP, indicating in CTR more robust functional connectivity within these states. In contrast, BNS 1 showed a higher variability in duration among FEP, suggesting altered temporal dynamics in this state. Transitions from BNS 5 to BNS 4 occurred more frequently in FEP.

**Figure 2.**
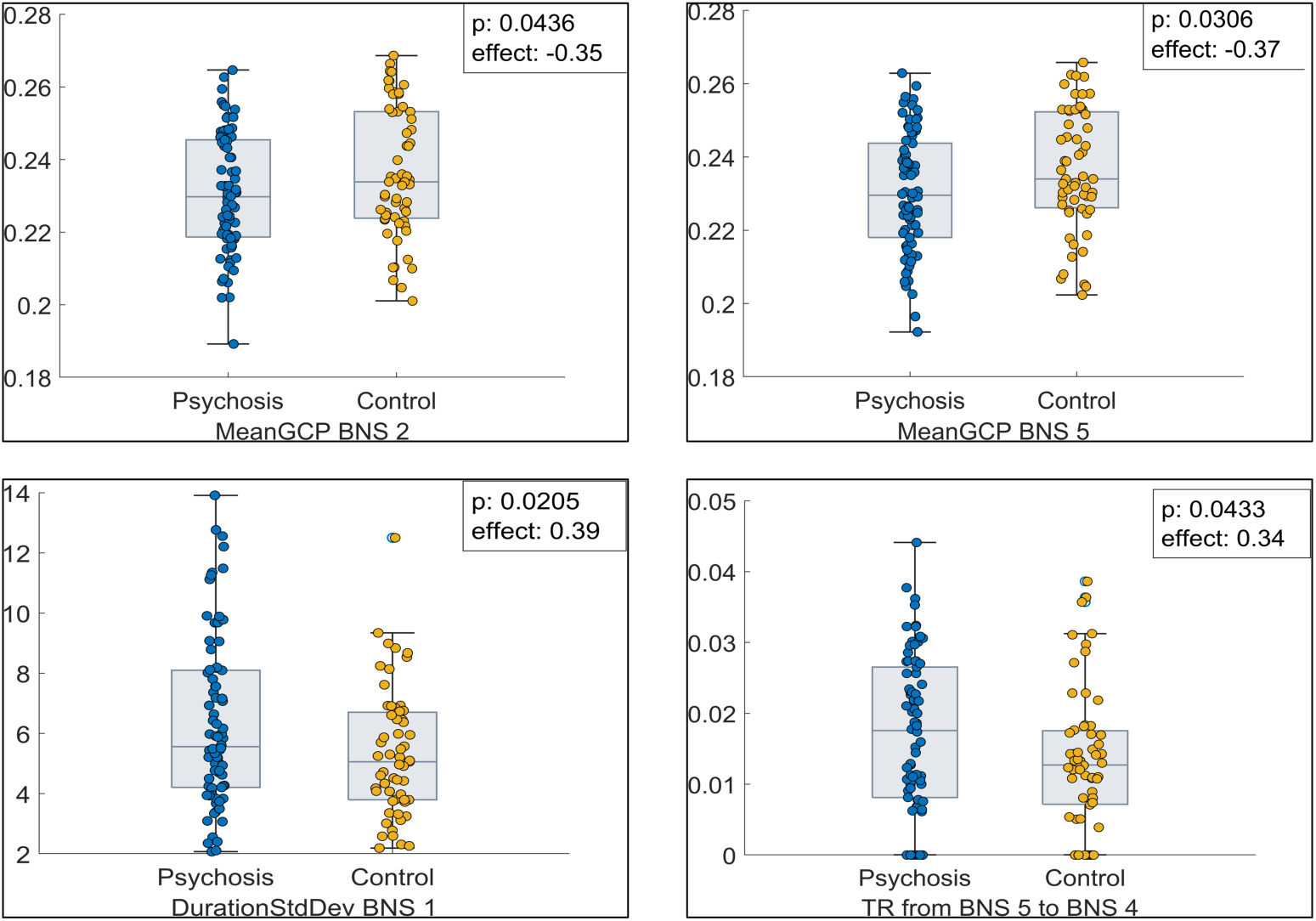
Boxplots with individual values of the significant differences (p<0.05) of the BNS metrics between FEP and CTR.

### 3. Correlation Analysis

Significant correlations were observed between BNS metrics and cognitive performance only in CTR, as highlighted in Figure 3.

**Figure 3.**
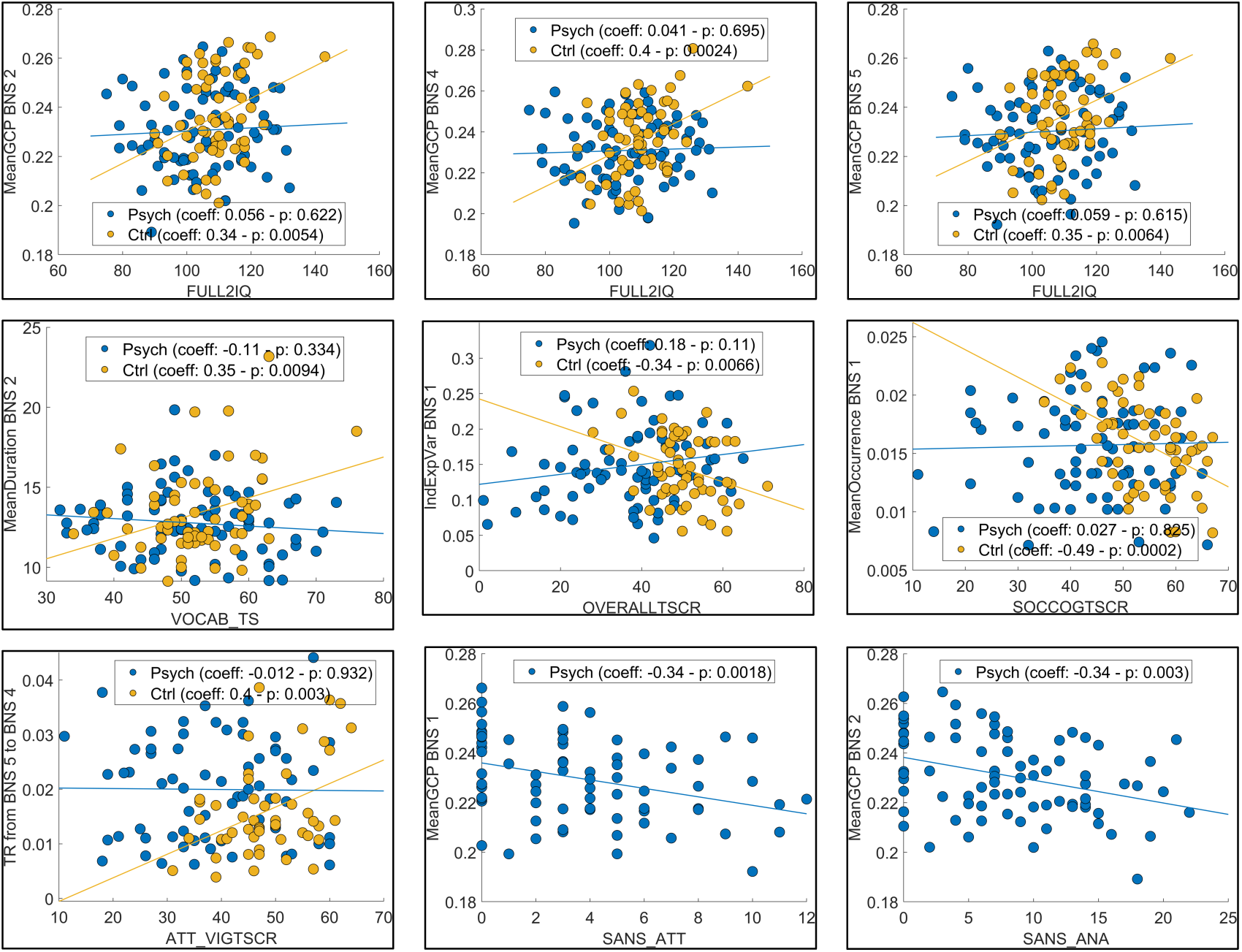
Scatterplots with regression lines of significant correlations (p<0.01) between the BNS metrics and the cognitive domains (measured by WASI and MCCB) in FEP (blu circles) and CTR (orange circles) and negative symptoms (assessed by SANS) in FEP.

In CTR, the metrics associated with the visually dominant BNS (BNS 2, 4, and 5) were positively correlated with the full IQ and Vocabulary scores on WASI. Furthermore, BNS 1 metrics were negatively correlated with social cognition and overall performance on the MCCB. Transition metrics, particularly from the right to the left hemisphere (BNS 5 to BNS 4), were positively correlated with attention and vigilance scores on the MCCB.

Conversely, in FEP, no significant correlations were found between BNS metrics and cognitive domain scores. However, negative correlations were identified between the GCP of BNS 1 and BNS 2 and the SANS’ Attention (SANS ATT) and Anhedonia–Asociality (SANS ANA) scales.

No significant correlations were observed between the BNS metrics and GAS.

## Discussion

The current study proposes a new methodology based on dynamic connectivity from EEG resting-state data and applies it to investigate neural correlates of psychosis and their relationship with cognitive and behavioral phenotypes. The clustering of alpha dynamic connectivity matrices revealed five distinct BNS, and some features showed significant differences between FEP and CTR. Particularly, BNS metrics correlated with cognition only in CTR and with negative symptoms in FEP, and, in line with the clinical characteristics of psychosis ^1–3,8^, this may underline potential neural correlates of the illness.

### 1. Interpretation of Identified BNS

Identifying distinct BNS provides crucial information on the neural dynamic processing at rest. To note, the five resulting BNS were observed in both groups without differences in terms of coverage and occurrence. This finding points out that our approach reveals brain connectivity reconfigurations in the alpha band during the resting state, a way for large-scale neural communication in the human brain without any specificity for psychotic illness.

BNS 2, 4, and 5, associated with visual regions, confirm a significant role for visual processing in brain network activity ^28^. This was expected, the recording paradigm being with the eyes open. However, the hemispheric differences observed in BNS 4 and BNS 5, with stronger left and right hemisphere activities, highlight brain function lateralization ^29^. BNS 1 and 3, marked by a higher representation of the default mode network (DMN), confirm the involvement of internally focused thought processes in the resting state ^30^.

### 2. Group Differences in BNS Metrics

The observed differences in GCP, duration variability, and transition probabilities between FEP and CTR are noteworthy. Compared to CTR, the lower GCP in BNSs 2 and 5 in FEP suggests a reduced alpha band functional connectivity. One of the possible explanations could be the instability of connections between ROIs in FEP. The greater variability in the duration of BNS 1 in FEP points to altered temporal dynamics, which may reflect a disruption in maintaining consistent brain states or the heterogeneity of psychosis ^7^. This extends the finding of altered MEG alpha neural dynamics previously observed in schizophrenia ^31^.

Additionally, the increased frequency of transitions from BNS 5 to BNS 4 in FEP suggests an altered pattern of neural communication between the right and left hemispheres compared to CTR.

Our results support data from other imaging modalities, which revealed group differences and altered network connectivity in schizophrenia ^32^.

### 3. Relationship between BNS Metrics and Clinical assessment

#### 3.1 Relationship Between BNS Metrics and Cognitive Performance

Contrary to our hypothesis, BNS features were only related to the cognitive domains in CTR but not FEP.

In CTR, the positive correlations between metrics of visually dominant BNS (2, 4, and 5) and cognitive performance, including IQ and vocabulary scores, indicate that efficient visual processing is linked to higher general cognitive abilities ^33^. The relation of BNS 4 (stronger activity on the left hemisphere) and BNS 5 (stronger on the right hemisphere) confirms the links between changes in lateralization and cognitive performance ^34^.

The negative correlations of BNS 1 metrics with social cognition and overall MCCB performance suggest that more pronounced DMN activity may burden these cognitive domains, confirming the complex links between this RSN and cognition ^35,36^. The transition metrics’ positive correlation with attention and vigilance scores implies that specific brain state transitions are crucial for sustained attention.

If, on the one hand, the relationship between descriptors of higher cognitive efficiency and BNS features in CTR suggests that BNS may serve as potential proxies for general brain functioning, on the other hand, the presence of specific associations indicates that BNS could be a viable candidate for characterizing the physiological processes underlying certain cognitive domains in CTR.

The lack of significant correlations in the psychosis group may be due to disrupted connectivity patterns that affect cognitive integration ^8^. As mentioned, this can also be explained by the heterogeneity of FEPs ^7^. A division into subgroups could be interesting to observe more specific discrimination between BNS metrics, cognitive performance, and clinical phenotype ^37–40^.

#### 3.2 Association Between BNS Metrics and Negative Symptoms

In line with our hypothesis, BNS features were associated only with negative symptoms and not with other psychopathological domains in FEP. The negative correlations between GCP in BNS 1 and BNS 2 with the SANS attention and anhedonia-asociality scales in FEP indicate that altered connectivity within these BNS is associated with specific psychopathological symptoms considered core of schizophrenia ^41,42^. These findings suggest that disruptions in particular brain states may underlie specific negative symptoms of psychosis, providing a potential pathophysiology and target for therapeutic interventions.

### 4. Advantages of the BNS approach

BNS metrics used in this study could discriminate patients from healthy subjects, revealing a strong relationship between the extracted metrics and cognitive profiles. They also correlated with the severity of negative symptoms. These findings are of interest, as they offer a new perspective in the analysis of psychosis. It points out that the methods and analysis of brain dynamics are crucial for clinical application as potential neural correlates of the disease.

From the resulting BNS sequence, on top of the current usual metrics derived from classical microstates ^43^, new metrics can be extracted based on complexity measures ^44^, which have already shown results in the observed dataset ^19^. Therefore, our approach can offer extensive metrics correlating well with cognitive and clinical scales, describing the brain dynamics in resting and potentially task-related states ^26,27^. These metrics could be input for future machine learning and classification work, which would help establish a patient-specific profile.

### 5. Methodological Considerations and Limitations

The current work focused on the alpha band based on previous literature. This could be extended to explore the behavior of other frequency bands, which could provide complementary but more speculative information.

Our study employs advanced source localization and clustering techniques, which offer a detailed analysis of brain network dynamics. However, several limitations should be noted. Substantial variability can be observed depending on the inverse solution used, the connectivity technique approach, and the clustering algorithm. Moreover, while advantageous for identifying common patterns, the group-level clustering approach may overlook individual variability. Additionally, potential confounding factors such as medication effects were not fully controlled, which could influence the findings. In future works, some steps of the methodology can be tuned to increase the granularity of our proposed approach: a more robust group and subgroup definition and individual-level *vs*. group-level clustering. Combining EEG with normative modeling is highly promising for developing patient-specific approaches ^45^.

### 6. Future Directions and Clinical Implications

The approach proposed in this study needs to be explored and validated with different datasets. Replicating our results and refining the methodology could lead to significant advancements in the field of psychosis. Like the classical microstate approach, defining BNS templates could facilitate easier categorization of resting-state brain activity and enable more global and comparable analyses between datasets. Future research should also consider longitudinal studies and multimodal neuroimaging approaches to validate and extend our findings. Investigating the stability of BNS over time and their relationship with treatment responses could provide valuable insights. The overall brain state field could be vital in addressing the relationship between neural dynamics, physiological changes, and phenotypes ^10^. Clinically, our findings have the potential to enhance diagnostic accuracy and guide the development of personalized interventions based on specific brain network disruptions in psychosis. Identifying reliable biomarkers of psychosis through EEG could significantly impact the management and treatment of this complex disorder, thereby improving patient outcomes.

## Methods and Materials

### 1- Participants

We examined rsEEG sourced from the OpenNeuro database ^46,47^ (data retrieved on November 29, 2023). The dataset comprised 78 FEP (23 female; age 22.9±4.8 years) and 60 age-matched CTR (26 female; 22.8±4.9). Table 1 contains a complete description with demographics and clinical and cognitive data.

**Table 1.** Dataset description. Continuous variables are reported in the format mean (range, standard deviation) and underwent a Welch’s t-test, and Hedges’g was computed. For the categorical variables, a χ^2^ test was performed.

|  | FEPs | Control | Test-statistic |
| --- | --- | --- | --- |
| **n** | 78 | 60 |  |
| Age (years) | 22.8 (13-36, 4.9) | 22.9 (12.8-38.3, 4.8) | $t_{136} = 0.07, p = 0.94, g = 0.01$ |
| Sex (F/M) | 23/55 | 26/34 | $\chi^2_1 = 2.84, p = 0.09$ |
| Race<br>(Asian/Black/Mixed/Undisclosed/White) | 6/25/6/1/40 | 13/6/2/0/39 | $\chi^2_4 = 15.15, p = 0.004$ |
| Ethnicity<br>(Hispanic/Not Hispanic/Unknown) | 3/75/0 | 5/54/1 | $\chi^2_2 = 2.62, p = 0.27$ |
| Medicated/Drug-Free/ Drug-Naïve | 51/14/13 | - |  |
| **WASI** |  |  |  |
| FULL IQ | 105 (75-132, 13.8) | 109 (90-143, 9.3) | $t_{136} = 2.03, p = 0.045, g = 0.34$ |
| MATRIX TS | 54 (32-68, 8.8) | 58.7 (45-70, 5.4) | $t_{136} = 3.82, p < 0.001, g = 0.63$ |
| VOCAB TS | 51.8 (32-73, 9.8) | 52 (34-76, 7.3) | $t_{136} = 0.2, p = 0.84, g = 0.03$ |
| **MCCB** |  |  |  |
| Attention/Vigilance | 38.4 (11-60, 11.8) | 47.3 (26-64, 8.5) | $t_{136} = 5.07, p < 0.001, g = 0.85$ |
| Speed of processing | 40.4 (8-74, 13.8) | 51.8 (35-73, 9.3) | $t_{136} = 5.67, p < 0.001, g = 0.96$ |
| Working memory | 41.9 (15-75, 12.3) | 48.2 (30-74, 8.9) | $t_{136} = 3.48, p < 0.001, g = 0.58$ |
| Verbal learning | 43.9 (25-67, 10.3) | 53.6 (37-77, 9) | $t_{136} = 5.82, p < 0.001, g = 0.99$ |
| Visual learning | 40.2 (8-63, 13) | 45.8 (25-61, 7.9) | $t_{136} = 3.13, p = 0.002, g = 0.52$ |
| Reasoning and problem-solving | 44.2 (18-63, 11.4) | 50.6 (33-65, 8.1) | $t_{136} = 3.81, p < 0.001, g = 0.64$ |
| Social cognition | 44.5 (11-66, 11.9) | 54.1 (35-67, 7.6) | $t_{136} = 5.71, p < 0.001, g = 0.95$ |
| Overall | 36.8 (1-65, 14.3) | 50.1 (28-71, 8) | $t_{136} = 6.89, p < 0.001,$<br>$g = 1.15$ |
| **SAPS** |  |  |  |
| Hallucinations | 6.2 (0-22, 5.9) | - |  |
| Delusions | 10.8 (0-29, 7.3) | - |  |
| Bizarre Behavior | 2.2 (0-17, 2.9) | - |  |
| Positive Formal Thought Disorder | 3.5 (0-20, 4.7) | - |  |
| **SANS** |  |  |  |
| Affective Flattening or Blunting | 8.9 (0-29, 7) | - |  |
| Alogia | 3.2 (0-12, 3.3) | - |  |
| Avolition-Apathy | 7 (0-14, 4) | - |  |
| Anhedonia-Asociality | 8.3 (0-22, 6) | - |  |
| Attention | 3.7 (0-12, 3.3) | - |  |
| **GAS** | 48.5 (23-90, 15) | - |  |

For each participant, a 5-minute rsEEG recording was acquired while they kept their eyes open. The EEG data were collected using the EEG channels of an Elekta Neuromag Vectorview MEG system. EEG was recorded using a low-impedance 10-10 system 60-channel cap at a sampling frequency of 1000 Hz. Since the data was available in 2 different EEG sensor arrays, a further validation analysis was performed, and the results are reported in the supplementary material (Table S1).

### 2 – Methods/Analysis

Figure 4 is a graphical abstract of our novel methodology that will be detailed in the following sections.

**Figure 4.**
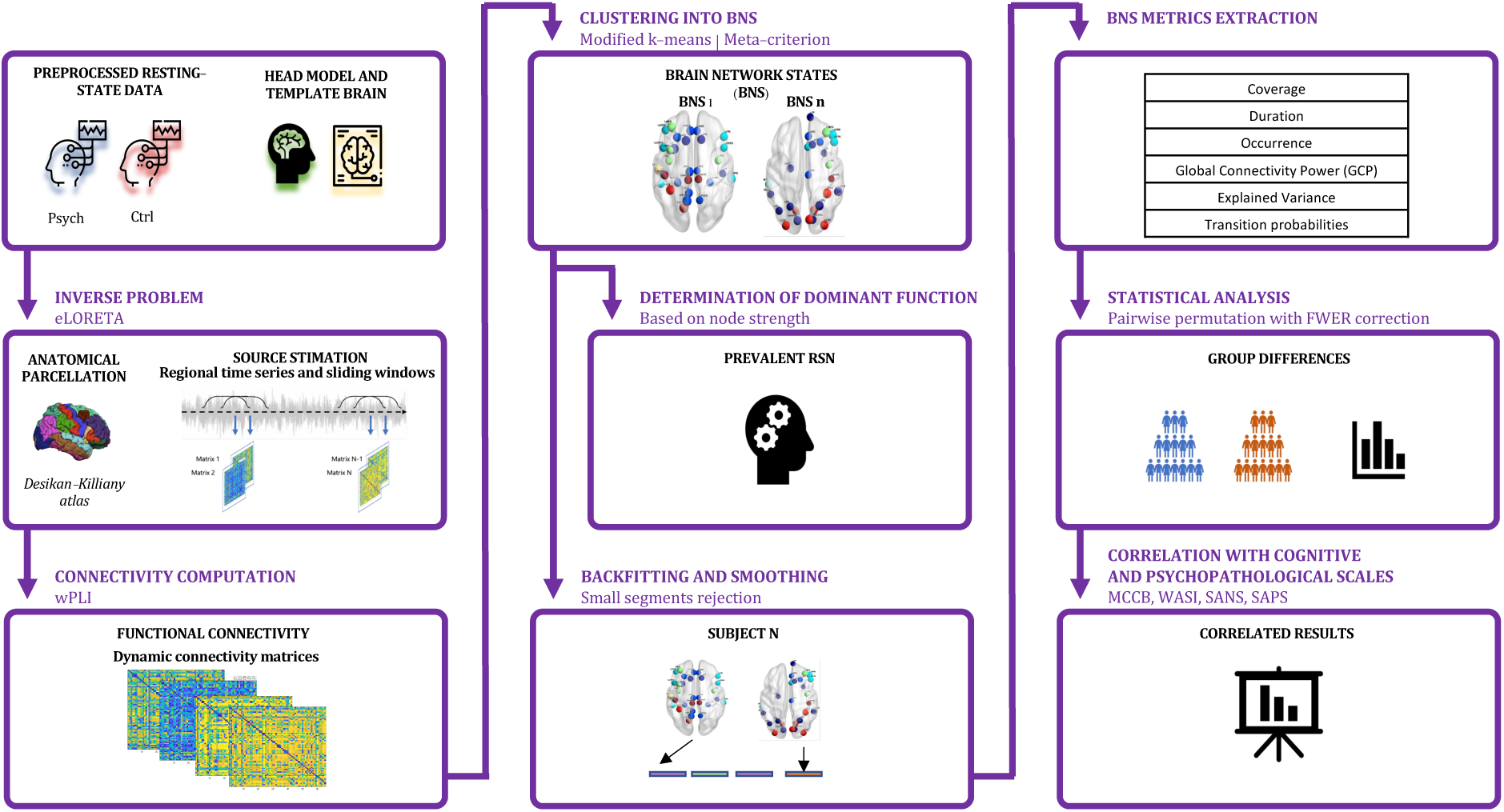
Graphical abstract of methods.

*2.1 - Preprocessing*

Pre-processing was performed with the EEGLAB toolbox ^48^.

First, a 1 Hz high-pass filter was applied. Then, a round of *low-level cleaning* to remove the artifacts was applied: flat channels were removed with a flatline criterion of 5 seconds, and line noise (60 Hz and harmonics) was removed. Channels with low correlation to their neighbors (< 0.6) were removed. The noisy channels were removed with a line noise criterion of 5. Artifact Subspace Reconstruction (ASR) was applied to correct non-periodic noise.

Later, *high-level cleaning* with ICA was performed using the PICARD algorithm ^49,50^. Independent components were labeled using the *ICLabel* procedure ^51^, and components classified as non-brain activity were rejected.

Removed channels were interpolated using the original channel locations using the spherical method. Data were re-referenced to the average reference.

After the initial preprocessing, a second passage of high-level cleaning was performed, including a second ICA decomposition, component labeling, and rejection to remove the residual periodic non-brain artifacts.

The final preprocessed data was used for further analysis.

#### 2.2- Source estimation and connectivity computation

We employed the exact Low-Resolution Brain Electromagnetic Tomography (eLORETA) algorithm for source localization ^52^. This method allowed us to estimate the cortical sources of EEG signals, focusing on 68 regions of interest (ROIs) as delineated by the Desikan-Killiany atlas ^53^. To perform this operation, we used the Boundary Element Method head model fitted to the ICBM MRI template, which is composed of three layers (scalp, outer skull, and inner skull), using the OpenMEEG ^54^ plugin from the Brainstorm toolbox ^55^. The ROI timeseries were estimated based on the average source strength of all voxels in each ROI.

Our primary interest lies in dynamic functional connectivity within the alpha frequency band (7-13 Hz), which aligns with recent studies showing alterations in the alpha band in EEG spectral ^15,16^ or connectivity ^18^ features. This was also based on MEG results, where part of the current dataset was analyzed ^19^, highlighting different networks in the alpha band.

We quantified connectivity patterns using the weighted phase lag index (wPLI), a measure robust to the effects of volume conduction and field spread ^56^. A sliding window approach (window size: 600ms, corresponding to 6 alpha frequency cycles as advised by Lachaux *et al*. ^57^, with a 50% overlap) was adopted to capture the temporal evolution of connectivity patterns, leading to 999 connectivity matrices per subject.

#### 2.3- BNS Clustering

Based on the principles of EEG microstate analysis, we conducted a clustering procedure using a modified k-means algorithm ^58,59^. This technique was chosen due to its ability to identify stable and recurring patterns of brain activity, as it was widely used in classical EEG microstate analyses ^43,60^, which we will refer to as BNS ^26,27^.

Unlike traditional approaches, our clustering was conducted at group rather than individual levels. The following pre-processing steps were performed on each subject’s data. The dynamic connectivity matrices, being symmetric, were vectorized by extracting only the lower part (resulting in 2278 elements for a 68×68 matrix), creating vectors sized 2278×999 for each subject. These vectors were then concatenated to generate a comprehensive dataset encompassing 138 subjects. This unified vector served as input for the clustering algorithm, following a methodology previously utilized by Duprez *et al*. (15).

Initially, we explored solutions iteratively within a range of 2 to 20, conducting 100 restarts with 1000 iterations each. To identify the optimal number of solutions, we designed a meta-criterion based on 11 clustering criteria ^59,61^, following an approach proposed by Custo *et al*. ^62^. The list of criteria and the meta-criterion computation are described in the supplementary material (Table S2). Based on the converging results from the several tests performed with this approach, we narrowed the interval to between 3 and 7 solutions, with 200 restarts and 1000 iterations, to address the computational weight.

#### 2.4- Smoothing and Backfitting

Following the identification of distinct BNS, we applied backfitting to each individual based on global map dissimilarity (GMD) ^63^, which is a distance measure based on the topography of each map and is signal invariant ^59^. Then, after the backfitting, we applied label smoothing based on small segment rejection ^59^: each BNS lasting less than five consecutive matrices was rejected and relabeled.

#### 2.5- BNS metrics and function

From these resulting datasets, we extracted the following BNS metrics: global connectivity power (GCP), i.e., the standard deviation of all connectivity indices, as derived from global field power (GFP) ^63^, explained variance (GEV), coverage, duration, occurrence, and transition probabilities ^43^. These metrics provided a comprehensive characterization of the BNS dynamics.

Additionally, we utilized node strength to link each BNS to resting-state networks (RSN) ^27^ and to characterize its prevalent functional role. Each node was assigned to one of seven RSNs (Figure 5) based on topographic criteria. We then normalized the strength of each RSN by the number of its nodes and computed the mean and standard deviation of the normalized values. The RSNs with a value above the mean + 1SD were defined as prevalent.

**Figure 5.**
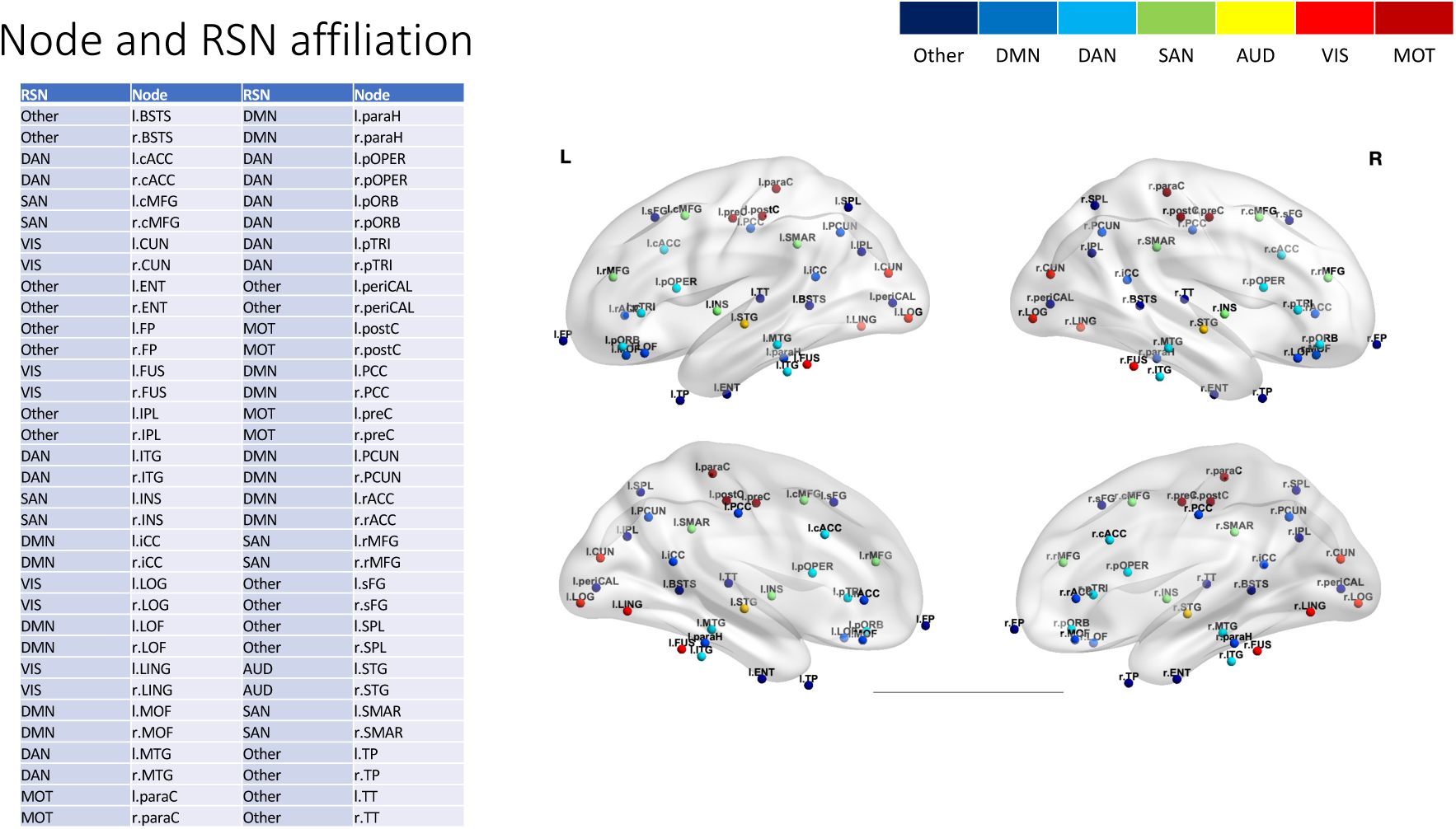
**List of ROIs and RSN**. AUD: Auditory Network; DAN: Dorsal Attentional Network; DMN: Default Mode Network; MOT: Motor Network; SAN: Salient Network; VIS: Visual Network

#### 2.6- Statistical analysis

A multivariate permutation approach was used for statistical analysis, with 10.000 permutations and family-wise error rate (FWER) correction based on max correction to control for multiple comparisons ^64^. To assess the differences in BNS metrics between FEP and CTR, we performed pairwise comparisons with a significant threshold set at α = 0.05. Furthermore, we conducted a Persons’ correlation analysis to explore the relationship between BNS metrics and cognitive performance, measured by the MATRICS Consensus Cognitive Battery (MCCB) and the Wechsler Abbreviated Scale of Intelligence (WASI), as well as psychopathological symptoms, assessed by the Scale for the Assessment of Positive Symptoms (SAPS), the Scale for the Assessment of Negative Symptoms (SANS), and the Global Assessment Scales (GAS). Due to the explorative nature of this analysis, only correlations with p-values lower than the 0.01 threshold are reported.

## Data availability

The raw data from this study is available from the OpenNeuro database.

## Acknowledgments & Disclosures

RA received grant support from the Department of Systems Medicine, University of Rome Tor Vergata, Italy (CUP I53C23001730007).

GDL was supported by #NEXTGENERATIONEU (NGEU) and funded by the Ministry of University and Research (MUR), National Recovery and Resilience Plan (NRRP), project MNESYS (PE0000006) – (DN. 1553 11.10.2022).

MH was supported by the ‘Region Bretagne’ – Inno R&D project n 23001155-.

The authors reported no biomedical financial interests or potential conflicts of interest.

The current manuscript has been posted on a preprint server: https://doi.org/10.1101/2024.06.04.597416

## Author Contribution

RA and GDL designed the analysis. RA conducted the main analysis presented and composed the figures and tables. RA and GDL prepared the first draft of the paper. MH, PG, and SS contributed to the project’s conceptualization, analysis plans, and paper preparation.

## Notes

### Competing Interest Statement

The authors have declared no competing interest.

### Summary of Updates

Results section updated Supplemental files updated Distribution/Reuse options updated Structure updated

## References

1. Maj, M. et al. The clinical characterization of the patient with primary psychosis aimed at personalization of management. World Psychiatry 20, 4–33 (2021).

2. Lieberman, J. A. & First, M. B. Psychotic Disorders. N. Engl. J. Med. 379, 270–280 (2018).

3. Jauhar, S., Johnstone, M. & McKenna, P. J. Schizophrenia. The Lancet 399, 473–486 (2022).

4. Voineskos, A. N. et al. Functional magnetic resonance imaging in schizophrenia: current evidence, methodological advances, limitations and future directions. World Psychiatry 23, 26– 51 (2024).

5. Gur, R. E., Keshavan, M. S. & Lawrie, S. M. Deconstructing Psychosis With Human Brain Imaging. Schizophr. Bull. 33, 921–931 (2007).

6. Scheepers, F. E., De Mul, J., Boer, F. & Hoogendijk, W. J. Psychosis as an Evolutionary Adaptive Mechanism to Changing Environments. Front. Psychiatry 9, 237 (2018).

7. Ganella, E. P. et al. Resting-state functional brain networks in first-episode psychosis: A 12- month follow-up study. Aust. N. Z. J. Psychiatry 52, 864–875 (2018).

8. McCutcheon, R. A., Reis Marques, T. & Howes, O. D. Schizophrenia—An Overview. JAMA Psychiatry 77, 201 (2020).

9. Biasiucci, A., Franceschiello, B. & Murray, M. M. Electroencephalography. Curr. Biol. 29, R80–R85 (2019).

10. Greene, A. S., Horien, C., Barson, D., Scheinost, D. & Constable, R. T. Why is everyone talking about brain state? Trends Neurosci. 46, 508–524 (2023).

11. Foster, M. & Scheinost, D. Brain states as wave-like motifs. Trends Cogn. Sci. 28, 492–503 (2024).

12. Wang, B., Zartaloudi, E., Linden, J. F. & Bramon, E. Neurophysiology in psychosis: The quest for disease biomarkers. Transl. Psychiatry 12, 100 (2022).

13. Sponheim, S. R., Clementz, B. A., Iacono, W. G. & Beiser, M. Clinical and biological concomitants of resting state EEG power abnormalities in schizophrenia. Biol. Psychiatry 48, 1088–1097 (2000).

14. Kam, J. W. Y., Bolbecker, A. R., O’Donnell, B. F., Hetrick, W. P. & Brenner, C. A. Resting state EEG power and coherence abnormalities in bipolar disorder and schizophrenia. J. Psychiatr. Res. 47, 1893–1901 (2013).

15. Clementz, B. A., Sponheim, S. R., Iacono, W. G. & Beiser, M. Resting EEG in first-episode schizophrenia patients, bipolar psychosis patients, and their first-degree relatives. Psychophysiology 31, 486–494 (1994).

16. Murphy, M. & Öngür, D. Decreased peak alpha frequency and impaired visual evoked potentials in first episode psychosis. NeuroImage Clin. 22, 101693 (2019).

17. Hassan, M. & Wendling, F. Electroencephalography Source Connectivity: Aiming for High Resolution of Brain Networks in Time and Space. IEEE Signal Process. Mag. 35, 81–96 (2018).

18. Di Lorenzo, G. et al. Altered resting-state EEG source functional connectivity in schizophrenia: the effect of illness duration. Front. Hum. Neurosci. 9, (2015).

19. Phalen, H., Coffman, B. A., Ghuman, A., Sejdić, E. & Salisbury, D. F. Non-negative Matrix Factorization Reveals Resting-State Cortical Alpha Network Abnormalities in the First-Episode Schizophrenia Spectrum. Biol. Psychiatry Cogn. Neurosci. Neuroimaging 5, 961–970 (2020).

20. Lottman, K. K. et al. Examining resting-state functional connectivity in first-episode schizophrenia with 7T fMRI and MEG. NeuroImage Clin. 24, 101959 (2019).

21. Kabbara, A., El Falou, W., Khalil, M., Wendling, F. & Hassan, M. The dynamic functional core network of the human brain at rest. Sci. Rep. 7, 2936 (2017).

22. O’Neill, G. C. et al. Dynamics of large-scale electrophysiological networks: A technical review. NeuroImage 180, 559–576 (2018).

23. Tabbal, J., Kabbara, A., Khalil, M., Benquet, P. & Hassan, M. Dynamics of task-related electrophysiological networks: a benchmarking study. NeuroImage 231, 117829 (2021).

24. Michel, C. M. & Koenig, T. EEG microstates as a tool for studying the temporal dynamics of whole-brain neuronal networks: A review. NeuroImage 180, 577–593 (2018).

25. Zanesco, A. P. Normative Temporal Dynamics of Resting EEG Microstates. Brain Topogr. 37, 243–264 (2024).

26. Duprez, J. et al. Spatio-temporal dynamics of large-scale electrophysiological networks during cognitive action control in healthy controls and Parkinson’s disease patients. NeuroImage 258, 119331 (2022).

27. Aubonnet, R. et al. Brain network dynamics in the alpha band during a complex postural control task. J. Neural Eng. 20, 026030 (2023).

28. Clayton, M. S., Yeung, N. & Cohen Kadosh, R. The many characters of visual alpha oscillations. Eur. J. Neurosci. 48, 2498–2508 (2018).

29. Güntürkün, O., Ströckens, F. & Ocklenburg, S. Brain Lateralization: A Comparative Perspective. Physiol. Rev. 100, 1019–1063 (2020).

30. Andrews-Hanna, J. R. The Brain’s Default Network and Its Adaptive Role in Internal Mentation. The Neuroscientist 18, 251–270 (2012).

31. Alamian, G. et al. Patient, interrupted: MEG oscillation dynamics reveal temporal dysconnectivity in schizophrenia. NeuroImage Clin. 28, 102485 (2020).

32. Houck, J. M. et al. Magnetoencephalographic and functional MRI connectomics in schizophrenia via intra- and inter-network connectivity. NeuroImage 145, 96–106 (2017).

33. Roelfsema, P. R. & De Lange, F. P. Early Visual Cortex as a Multiscale Cognitive Blackboard. Annu. Rev. Vis. Sci. 2, 131–151 (2016).

34. Wu, X. et al. Dynamic changes in brain lateralization correlate with human cognitive performance. PLOS Biol. 20, e3001560 (2022).

35. Smith, V., Mitchell, D. J. & Duncan, J. Role of the Default Mode Network in Cognitive Transitions. Cereb. Cortex 28, 3685–3696 (2018).

36. Smallwood, J. et al. The default mode network in cognition: a topographical perspective. Nat. Rev. Neurosci. 22, 503–513 (2021).

37. Clementz, B. A. et al. Testing Psychosis Phenotypes From Bipolar–Schizophrenia Network for Intermediate Phenotypes for Clinical Application: Biotype Characteristics and Targets. Biol. Psychiatry Cogn. Neurosci. Neuroimaging 5, 808–818 (2020).

38. Clementz, B. A. et al. Identification of Distinct Psychosis Biotypes Using Brain-Based Biomarkers. Am. J. Psychiatry 173, 373–384 (2016).

39. Clementz, B. A. et al. Psychosis Biotypes: Replication and Validation from the B-SNIP Consortium. Schizophr. Bull. 48, 56–68 (2022).

40. Tang, S. X. et al. Functional phenotypes in schizophrenia spectrum disorders: defining the constructs and identifying biopsychosocial correlates using data-driven methods. Schizophrenia 10, 58 (2024).

41. Galderisi, S., Mucci, A., Buchanan, R. W. & Arango, C. Negative symptoms of schizophrenia: new developments and unanswered research questions. Lancet Psychiatry 5, 664–677 (2018).

42. Bègue, I., Kaiser, S. & Kirschner, M. Pathophysiology of negative symptom dimensions of schizophrenia – Current developments and implications for treatment. Neurosci. Biobehav. Rev. 116, 74–88 (2020).

43. Nagabhushan Kalburgi, S., et al. MICROSTATELAB: The EEGLAB Toolbox for Resting-State Microstate Analysis. Brain Topogr. (2023) doi:10.1007/s10548-023-01003-5.

44. Von Wegner, F. et al. Complexity Measures for EEG Microstate Sequences: Concepts and Algorithms. Brain Topogr. 37, 296–311 (2024).

45. Ebadi, A. et al. Beyond homogeneity: Charting the landscape of heterogeneity in psychiatric electroencephalography. Preprint at 10.1101/2024.03.04.583393 (2024).

46. 46. Salisbury, D., Seebold, D. & Coffman, B. EEG: First Episode Psychosis vs. Control Resting Task 1. [object Object] 10.18112/OPENNEURO.DS003944.V1.0.1 (2022).

47. 47. Salisbury, D., Seebold, D. & Coffman, B. EEG: First Episode Psychosis vs. Control Resting Task 2. [object Object] 10.18112/OPENNEURO.DS003947.V1.0.1 (2022).

48. Delorme, A. & Makeig, S. EEGLAB: an open source toolbox for analysis of single-trial EEG dynamics including independent component analysis. J. Neurosci. Methods 134, 9–21 (2004).

49. Ablin, P., Cardoso, J.-F. & Gramfort, A. Faster Independent Component Analysis by Preconditioning With Hessian Approximations. IEEE Trans. Signal Process. 66, 4040–4049 (2018).

50. 50. Frank, G., Makeig, S. & Delorme, A. A Framework to Evaluate Independent Component Analysis applied to EEG signal: testing on the Picard algorithm. Preprint at 10.48550/ARXIV.2210.08409 (2022).

51. Pion-Tonachini, L., Kreutz-Delgado, K. & Makeig, S. The ICLabel dataset of electroencephalographic (EEG) independent component (IC) features. Data Brief 25, 104101 (2019).

52. Pascual-Marqui, R. D. et al. Assessing interactions in the brain with exact low-resolution electromagnetic tomography. Philos. Trans. R. Soc. Math. Phys. Eng. Sci. 369, 3768–3784 (2011).

53. Desikan, R. S. et al. An automated labeling system for subdividing the human cerebral cortex on MRI scans into gyral based regions of interest. NeuroImage 31, 968–980 (2006).

54. Gramfort, A., Papadopoulo, T., Olivi, E. & Clerc, M. OpenMEEG: opensource software for quasistatic bioelectromagnetics. Biomed. Eng. OnLine 9, 45 (2010).

55. Tadel, F., Baillet, S., Mosher, J. C., Pantazis, D. & Leahy, R. M. Brainstorm: A User-Friendly Application for MEG/EEG Analysis. Comput. Intell. Neurosci. 2011, 1–13 (2011).

56. Vinck, M., Oostenveld, R., Van Wingerden, M., Battaglia, F. & Pennartz, C. M. A. An improved index of phase-synchronization for electrophysiological data in the presence of volume-conduction, noise and sample-size bias. NeuroImage 55, 1548–1565 (2011).

57. Lachaux, J.-P. et al. STUDYING SINGLE-TRIALS OF PHASE SYNCHRONOUS ACTIVITY IN THE BRAIN. Int. J. Bifurc. Chaos 10, 2429–2439 (2000).

58. Pascual-Marqui, R. D., Michel, C. M. & Lehmann, D. Segmentation of brain electrical activity into microstates: model estimation and validation. IEEE Trans. Biomed. Eng. 42, 658–665 (1995).

59. Poulsen, A. T., Pedroni, A., Langer, N. & Hansen, L. K. Microstate EEGlab toolbox: An introductory guide. Preprint at 10.1101/289850 (2018).

60. Tarailis, P., Koenig, T., Michel, C. M. & Griškova-Bulanova, I. The Functional Aspects of Resting EEG Microstates: A Systematic Review. Brain Topogr. 37, 181–217 (2024).

61. José-García, A. & Gómez-Flores, W. CVIK: A Matlab-based cluster validity index toolbox for automatic data clustering. SoftwareX 22, 101359 (2023).

62. Custo, A. et al. Electroencephalographic Resting-State Networks: Source Localization of Microstates. Brain Connect. 7, 671–682 (2017).

63. Murray, M. M., Brunet, D. & Michel, C. M. Topographic ERP Analyses: A Step-by-Step Tutorial Review. Brain Topogr. 20, 249–264 (2008).

64. Crosse, M. J., Foxe, J. J. & Molholm, S. PERMUTOOLS: A MATLAB Package for Multivariate Permutation Testing. (2024) doi:10.48550/ARXIV.2401.09401.

