## Supplementary material for "Resting-State Electroencephalography Alpha Dynamic Connectivity: Quantifying Brain Network State Evolution in Individuals with Psychosis"

### Table S1. Influence of channel array on the ROI spectral activity.

The sample set combines two sets recorded with two different EEG channel arrays from an Elekta Neuromag Vectorview system, using a low-impedance 10-10 system 60-channel cap. To confirm using those two sets in the same study, we ran a repeated measures ANOVA based on the ROIs' absolute power spectral density (Welch's method) in the alpha band (7-13Hz).

The results show a p-value higher than the 0.05 threshold, highlighting that the montage did not influence the ROI estimation. The two montages can be found on the OpenNeuro database [1,2] and referenced in the Methods section of the main text.

**Between Subjects Effects**

| Cases | Sum of Squares | df | Mean Square | F | p |
|---|---|---|---|---|---|
| type | $3.737\times10^{-17}$ | 1 | $3.737\times10^{-17}$ | 5.920 | 0.016 |
| Array | $1.062\times10^{-17}$ | 1 | $1.062\times10^{-17}$ | 1.682 | 0.197 |
| type × Array | $2.020\times10^{-17}$ | 1 | $2.020\times10^{-17}$ | 3.200 | 0.076 |
| Residuals | $8.460\times10^{-16}$ | 134 | $6.313\times10^{-18}$ | | |

*Note.* Type III Sum of Squares

**Table S2 – List of clustering criteria for the Meta-criterion**

| *Criterion* |
| --- |
| Krzanowski-Lai criterion |
| Normalised Krzanowski-Lai criterion |
| Cross-validation criterion |
| Calinski-Harabasz index |
| Enhanced Davies-Bouldin index |
| PBM index |
| S_Dbw validity index |
| Score Function index |
| SV index |
| Xie-Beni index |
| WB index |

**Merging the 11 Criteria**

The following process combined the 11 criteria into a single overarching metric.

In the first step, each of the 11 criteria was ranked separately by sorting the values from lowest to highest and using the relative positions.

Then, the criteria were merged using a formula designed to achieve two key goals: maximize average ranking and favor unanimity - having all 11 criteria in close agreement. To do so, maximizing the signal-to-noise ratio across criteria is defined as achieving maximum unanimity.

The meta-criterion equations can be found in the supplementary material of *Custo et al.* [3].

**References**


1. **Salisbury, D., Seebold, D. & Coffman, B. EEG: First Episode Psychosis vs. Control Resting Task 1. [object Object] https://doi.org/10.18112/OPENNEURO.DS003944.V1.0.1 (2022).**
2. **Salisbury, D., Seebold, D. & Coffman, B. EEG: First Episode Psychosis vs. Control Resting Task 2. [object Object] https://doi.org/10.18112/OPENNEURO.DS003947.V1.0.1 (2022).**
3. **Custo, A. *et al.* Electroencephalographic Resting-State Networks: Source Localization of Microstates. *Brain Connect.* 7, 671–682 (2017).**
